# The relative fitness of drug resistant *Mycobacterium tuberculosis*: a modelling study of household transmission in Peru

**DOI:** 10.1101/195313

**Authors:** Gwenan M. Knight, Mirko Zimic, Sebastian Funk, Robert H. Gilman, Jon S. Friedland, Louis Grandjean

## Abstract

The relative fitness of drug resistant versus susceptible bacteria in an environment dictates resistance prevalence. Estimates for the relative fitness of resistant *Mycobacterium tuberculosis* (*Mtb*) strains are highly heterogeneous and mostly derived from *in-vitro* experiments. Measuring fitness in the field allows us to determine how the environment influences resistance spread.

We designed a household structured, stochastic mathematical model to estimate the fitness costs associated with multi-drug resistance (MDR) carriage in *Mtb* in Lima, Peru between 2010-2013. By fitting the model to data from a large prospective cohort study of TB disease in household contacts we estimated the fitness, relative to susceptible strains with a fitness of 1, of MDR-*Mtb* to be 0.33 (95% credible interval: 0.17-0.54) or 0.39 (0.26-0.58), if only transmission or progression to disease, respectively, was affected. The relative fitness of MDR-*Mtb* increased to 0.57 (0.43-0.73) when the fitness cost influenced both transmission and progression to disease equally.

We found the average relative fitness of MDR-*Mtb* circulating within households in Lima, Peru between 2010-2013 to be significantly lower than concurrent susceptible-*Mtb*. If these fitness levels do not change, then existing TB control programmes are likely to keep MDR-TB prevalence at current levels in Lima, Peru.

## Background

*Mycobacterium tuberculosis* (*Mtb*) is a highly prevalent bacterium, thought to infect just under a quarter of the wrld’s population [1]. Treatment of tuberculosis (TB) disease is not simple and drug-susceptible tuberculosis (DS-TB) requires a multiple drug regimen taken for at least 6 months [2]. Multidrug-resistant tuberculosis (MDR-TB) treatment regimens are significantly longer, cause serious side effects and are very expensive [3]. Whilst currently 5% of all TB cases globally are estimated to be MDR-TB [2], predicting the future burden of DS- and MDR-TB is essential for TB control programmes.

One key parameter that determines the future prevalence of drug resistant TB is the relative fitness of drug resistant *Mtb* strains as compared to drug susceptible *Mtb* strains [4-7]. Fitness is a complex, environment-dependent trait that can be defined as the ability of a pathogen to survive, reproduce, be transmitted and cause secondary cases of disease. These abilities are affected by multiple environmental factors such as a host’s genetics, the current TB treatment regimen and other risk factors for transmission, which are all time-varying. The importance of this parameter has been highlighted by several mathematical models which show how even small changes in its value can predict widely varying future levels of MDR-TB burden [4-6, 8, 9]. Thus, gaining environment dependent, accurate estimates of fitness is of critical importance.

Within *Mtb*, it has been shown that the appearance of drug resistance mutations affects fitness [10-12]. These previous studies have shown that resistant *Mtb* is, usually, less fit than susceptible *Mtb* under a range of fitness definitions: either by demonstrating a lower growth rate *in vitro* (e.g. [13]), less progression to disease after inoculation in guinea pigs (e.g. [14]) or a lower chance of causing secondary cases of disease (e.g. [12, 15]). The latter definition is important for epidemiological predictions of burden, whilst the first provides the potential underlying biological cause. The epidemiological fitness of a *Mtb* strain can be split into an ability to (1) cause secondary infections (transmission) and (2) cause subsequent active disease (progression). For example, resistant *Mtb* may be transmitted equally as well, but subsequent disease rates in those infected may be lower or less severe. For *Mtb* this split is especially pertinent due to the importance of the latent, non-infectious, stage of disease.

Also highly important for *Mtb* is the spatial location of transmission [16]. Few studies have considered the critical influence of household structure on transmission of *Mtb*. To our knowledge, no studies have considered the spread of drug-resistant tuberculosis in the context of a household-structured stochastic mathematical model.

The difference in definitions of fitness and corresponding experimental data makes translation from data analysis to predictive mathematical modelling difficult. Here, we tackle this problem by fitting a mathematical model to a detailed data set on the transmission of *Mtb* strains collected in a large cohort study of households undertaken in Lima, Peru between 2010 and 2013 [17]. We derive estimates of fitness in this specific setting with different fitness definitions (either effects on transmission and/or progression to disease) and test the robustness of these estimates under a range of assumptions. These parameters will allow for better predictions of future MDR-TB levels and an improved understanding of MDR-TB spread.

## Methods

### Data

The details of the study and participants can be found in [17]. Briefly, 213 and 487 households were recruited with an index case of diagnosed MDR- or DS-TB respectively during 2010 to 2013. Households were followed up for variable periods of time up to a maximum of 3 years (Figure S1). During the study households were visited every 6 months, and household contacts were monitored for TB disease. It was found that 35/1055 (3.32%, 95% CI [2.32, .4.58]), of the MDR-TB contacts, and 114/2356 (4.84%, 95% CI [4.01, 5.78]), of the DS-TB contacts developed TB disease, suggesting that DS-TB has higher fitness. There were no significant differences between cohorts by HIV status, age, gender or household size [17].

The specific data used to calibrate the model was 1) the incidence of MDR-TB and 2) DS-TB in households with an index DS-TB case and 3) the incidence of MDR-TB and 4) DS-TB in households with an index MDR-TB case (Table 1). The percentages of incident cases with resistance profiles matching the index was used to multiply the incidence levels accordingly.

**Table 1:**
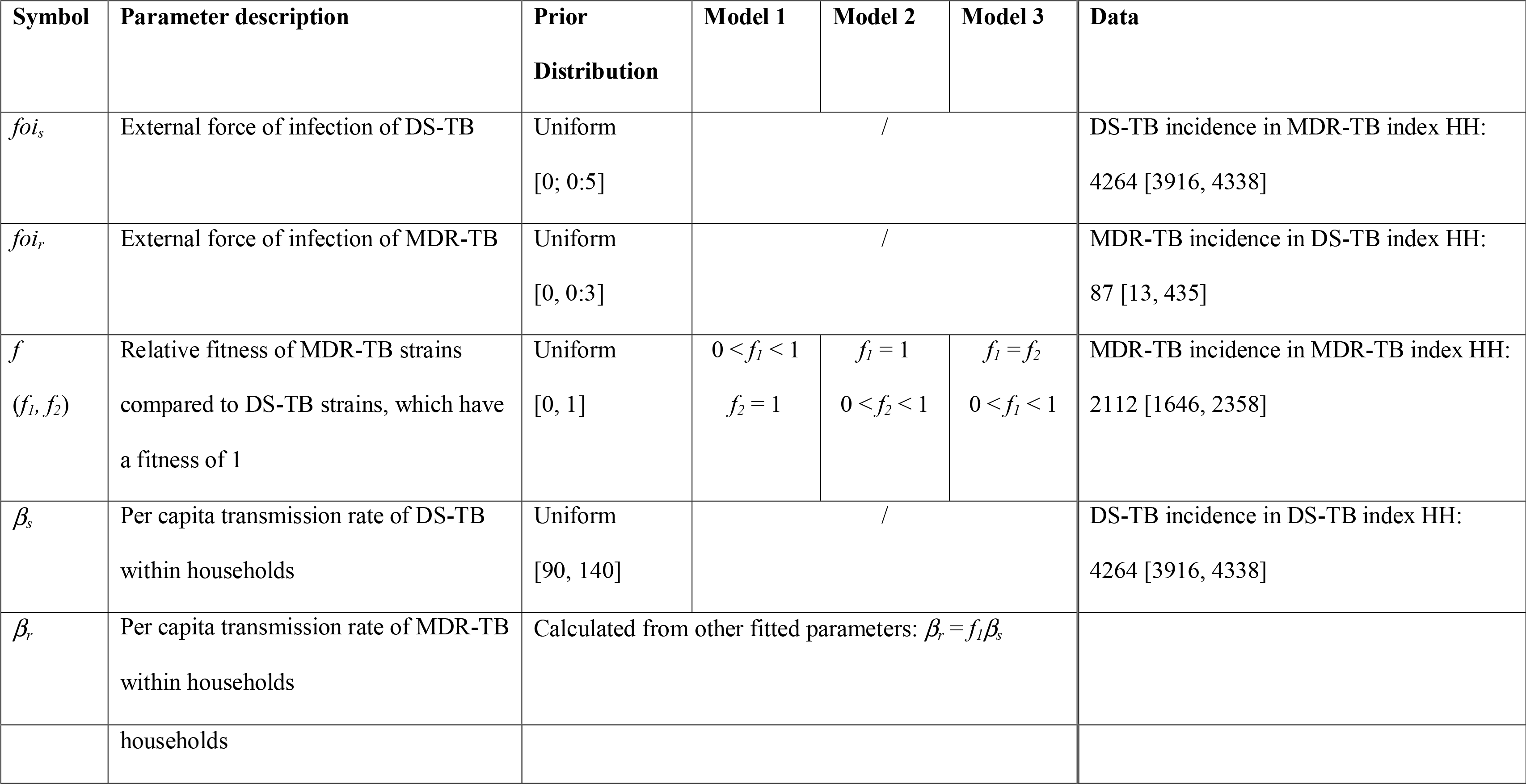
Fitted parameters with description, prior distributions, any differences by model structure and data used for fitting. All parameters are fitted to the TB incidence date from the household (HH) study (17). The three models have different assumptions around the effect of decreased fitness, with *f* varying to be *f_1_* (affects transmission rate) or *f_2_* (affects progression to disease rate) (see Figure 1).

### Model structure

The mathematical model was a standard two-strain dynamic TB model (Figure 1), with transmission modelled at the level of the household. A Gillespie stochastic simulation algorithm in *R* [18] was developed using the R package “GillespieSSA” [19]. Using a stochastic transmission model was important as the model was implemented independently in households where the small populations mean stochastic effects are highly important. We assumed that saturation of transmission could occur and hence scaled our transmission rate by the size of the household (number of people), assuming households have the same ventilation level (or at least that this did not vary by index case *Mtb* resistance status) and within-household homogeneous mixing [20]. This assumption of frequency-dependent transmission means that in households with more people, household members are assumed to have lower individual chance of infection from an active disease case than in smaller households, due to decreased exposure. This has been observed for another airborne pathogen, influenza [21] and was explored in sensitivity analysis where we also considered density-dependent transmission. All natural history parameters were taken from the literature, are listed in Table 2 and the dynamics explained in the legend to Figure 1.

**Table 2:** Parameter values with description and baseline values. All prior distributions were uniform.

| Symbol | Parameter description | Baseline value | Prior distribution | Notes and references |
| --- | --- | --- | --- | --- |
| $Nr$ | Number of households with MDR-TB index case | 213 | / | [24] |
| $Ns$ | Number of households with DS-TB index case | 487 | / | [24] |
| $h$ | Household size | 2 - 15 | / | [17] |
| $p$ | Proportion of (re-)infected individuals which progress to the “latent fast” state | 0.15 | 0.08-0.25 | [38-40] |
| $\chi$ | Protection from developing active TB upon re-infection | 0.35 | 0.25-0.45 | [38, 41-44] |
| $\phi$ | Rate of reactivation among those latently infected per year | $1.13 \times 10^{-4}$ | $1 - 3 \times 10^{-4}$ | [38, 41, 42, 44-46] |
| $\varepsilon$ | Probability of acquiring new drug resistance during treatment | 0.008 | 0.005-0.01 | [47] |
| $d$ | Proportion of new active cases which directly become infectious | 0.5 | 0.25-0.75 | [38, 44, 48, 49] |
| $\mu$ | Background death rate | $1/77 = 0.013$ | 0.012-0.014 | Inverse of average life expectancy in Peru [50] |
| $\mu_A$ | Additional death rate of those actively infected and infectious per year | 0.26 | 0.2-0.4 | [38] |
| $n$ | Annual rate of natural cure for | 0.2 | 0.15-0.25 | [38] |
|  | TB cases (returns to latent state) |  |  |  |
| $\omega_s$ | Proportion of DS-TB active cases detected and treated per year | 0.8; 2 | 0.5-0.95 | For 2012 [2] for prestudy; In study: screen every 6 months |
| $\omega_r$ | Proportion of MDR-TB active cases detected and treated per year | 0.64; 2 | 0.2-0.9 | 79% of the above 80% ($\omega_s$) found that received DST in 2012 [51]; In study: screen every 6 months |
| $(1 - k_s)$ | Proportion of DS-TB active cases started on treatment that are successfully cured | 0.74 | 0.5-0.9 | [51, 52] (for midpoint of study) |
| $(1 - k_r)$ | Proportion of MDR-TB active cases started on treatment that are successfully cured | 0.6 | 0.2-0.9 | For 2012 [2] |
| $p_f$ | Progression rate of latent fast individuals to active disease | 0.2 | 0.1-0.9 | Duration of fast latency period of 5 years [42] |

**Figure 1:**
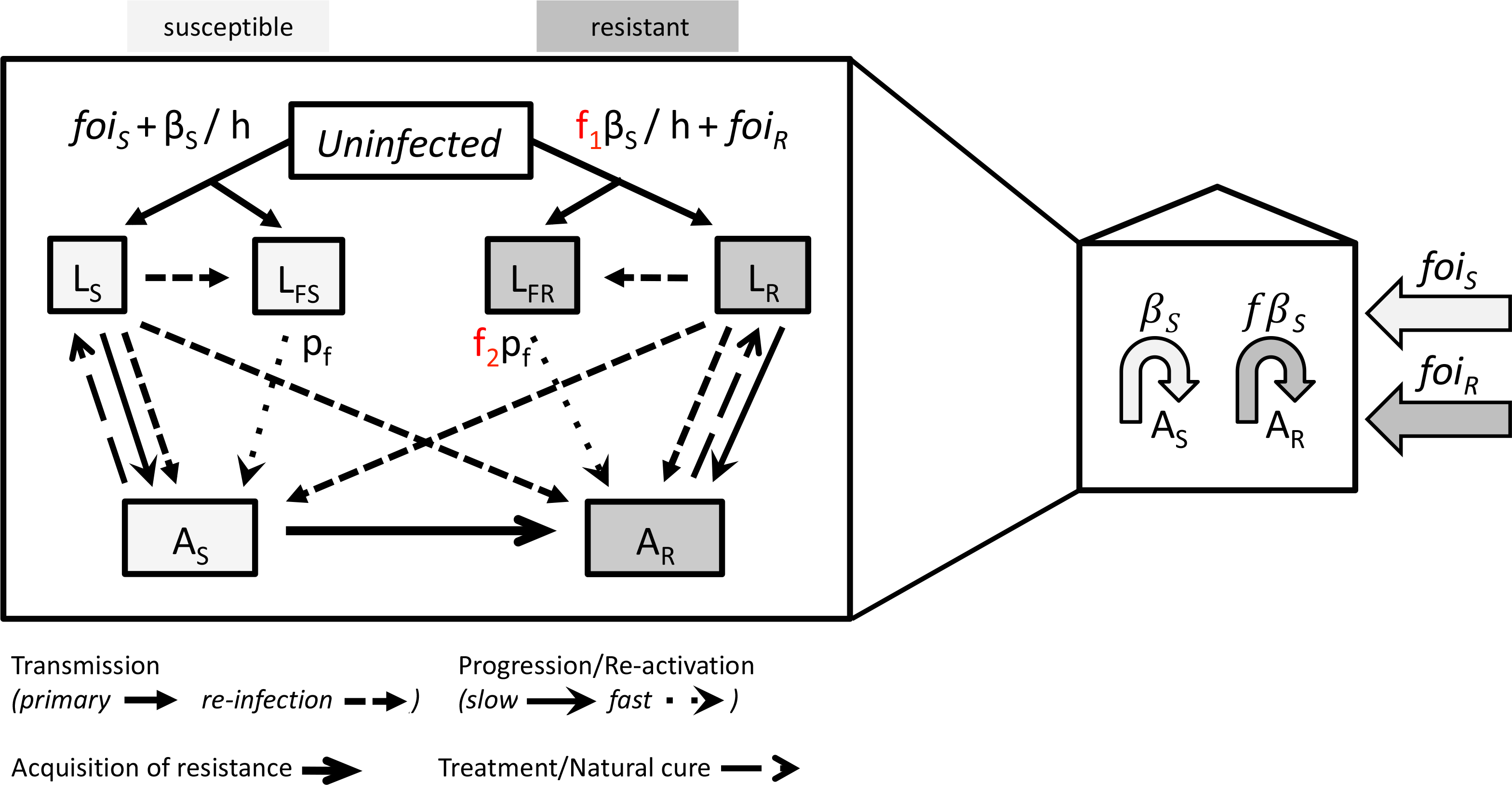
A standard natural history, transmission model for two strains (susceptible and resistant) of *Mtb* was used (diagram on the left). Uninfected people become infected at a rate dependent on the number of active cases (dynamic transmission). Once infected, the majority of people (85%) are assumed to enter a Latent slow (LS / LR) state. The remainder enter a rapid progression (Latent fast, LFS / LFR) state which has a higher rate of progression to active disease (AS / AR). Resistance mutations are acquired during active disease. Those with active disease recover to the Latent slow state via treatment or natural cure. The fitness cost to resistance is assumed to affect the rate of transmission (*f_1_*) or the rate at which those latently infected with MDR-TB progress to active disease (*f_2_*). Only the effect on primary transmission of *f_1_* is highlighted here, but reinfection is also decreased. *f_1_* and *f_2_* are set at 1 or allowed to vary between 0 and 1 in the three separate models: *f_1_* in Model 1, *f_2_* in Model 2 and both *f_1_* and *f_2_* in Model 3. The four estimated parameters (shown in the diagram on the right) were rates of internal transmission (*β_s_*, *f*) and the external forces of infection (*foi_s_* + *foi_R_*).

Four parameters were estimated from the data (Table 1, Figure 1): (1) the per capita transmission rate of DS-TB within households (*β*_S_), (2) the relative fitness of MDR-*Mtb* strains vs. DS-*Mtb* strains (*f*) expressed as an effect on transmission or progression or both, and the external (to households) force of infection (*foi*) of (3) DS-TB *foi*_s_ and (4) MDR-TB *foi*_r_.

Our main outcome was the impact of resistance on transmission rates, but we also explored the impact on an approximate effective reproduction number (see S1 text).

### Three model formulations

Resistant strains were allowed to have an equal or lower fitness relative to susceptible strains. The mechanisms behind this reduction were estimated to affect two different rates: the transmission rate, the rate of progression to disease, or both (Figure 1). We assumed that the fitness of the resistant strains could not rise above that of susceptible strains due to the data from the household cohort [17]. Model 1 (transmission fitness cost model) assumed that fitness costs directly affected the number of secondary infections by reducing the transmission parameter for MDR-*Mtb* (0 < *f_1_ < 1, f_2_* = 1, Figure 1). This is the standard assumption for the effect of resistance on fitness for transmission dynamic models of *Mtb* [6, 9, 22] and other pathogens [23]. Model 2 (progression fitness cost model) assumed that although MDR-TB transmission occurred at the same rate as DS-TB, there is a fitness cost to progression to disease (*f_1_* =1, *0 < f_2_* < 1, Figure 1). Model 3 assumed that there was a fitness cost to both transmission and progression, and that the cost was the same for both processes (0 < *f_1_* = *f_2_* < 1, Figure 1). We could not explore a Model with fitness affecting both processes at differing levels as we did not have data on levels of infection. Without this data, a model with high transmission fitness cost but low progression cost would be equally as likely as a model with a low transmission fitness cost but a high progression cost and hence would be uninformative. Note that fixing either *f_1_* or *f_2_* equal to one is the same as ignoring this parameter altogether and leaving the multiplied rate at its background level as they are both scalar constant parameters with no units.

### Model simulation

The model initially sampled 700 household sizes (with replacement) from the exact distribution of household sizes in the trial [24]. Initial number with latent infection were sampled from a normal distribution generated by data from the literature [2, 25] (S1 text). The model was then simulated for 10 years with a MCMC sampled set of the four unknown parameters (pre-study period), capturing transmission within the household prior to enrolment in the household study. A random time point from over this 10-year period in which there was at least one active case with the same sensitivity as the initial case in the household (i.e. DS-TB or MDR-TB) was taken to be the time the household entered the study and the active index case was detected. This allows for simulation of changes in latency in the household and provides initial conditions dependent upon each parameter sample.

The above randomly chosen time point of entry to the study was taken to be the initial conditions for the simulation of the model that was fitted to the household study [17] (study period). The same values of the four unknown parameters was used as in the pre-study period and the simulation time for each household was randomly sampled (with replacement) from the exact distribution of follow-up times in the study (Figure S1). The only parameter that changed, to match the altered patient care in the study, was the case detection rate which increased for the study period from the WHO estimates to a screen occurring every 6 months (Table 2).

The TB incidence from the model was calculated by determining the total number of new cases of active TB in all 700 households over the follow-up time, and dividing this by the total number of follow-up years in these households. The total number of follow-up years was a product of the number of household members and the follow-up time for the household taking into account any deaths over this time. We assumed that no-one left the households other than by death (natural or due to TB). For a detailed overview of the process see Figure S2.

### Model fitting

Approximate Bayesian Computation (ABC) was paired with Markov chain Monte Carlo (MCMC) methods to estimate the four unknown parameters [26]. All other parameters were kept fixed at their baseline value (Table 2). The summary statistic used was the TB incidence from the model falling within the 95% CI for all four TB incidence measures from the data. Uniform priors were assumed for all four parameters (Table 1).

To estimate the standard deviation required for the MCMC for the four unknown parameters, Latin Hypercube Sampling (LHS) from the prior ranges was initially used (Stage A, S1 text). The empirical standard deviation from the accepted fits was then used as proposal distribution of a Metropolis-Hastings MCMC sampler (Stage B), used to estimate posterior probabilities of the parameters.

We used the generated trajectories to consider the probability of remaining free of tuberculosis from the model output and compare the general trends to the data (Figure 2 from [17]).

**Figure 2:**
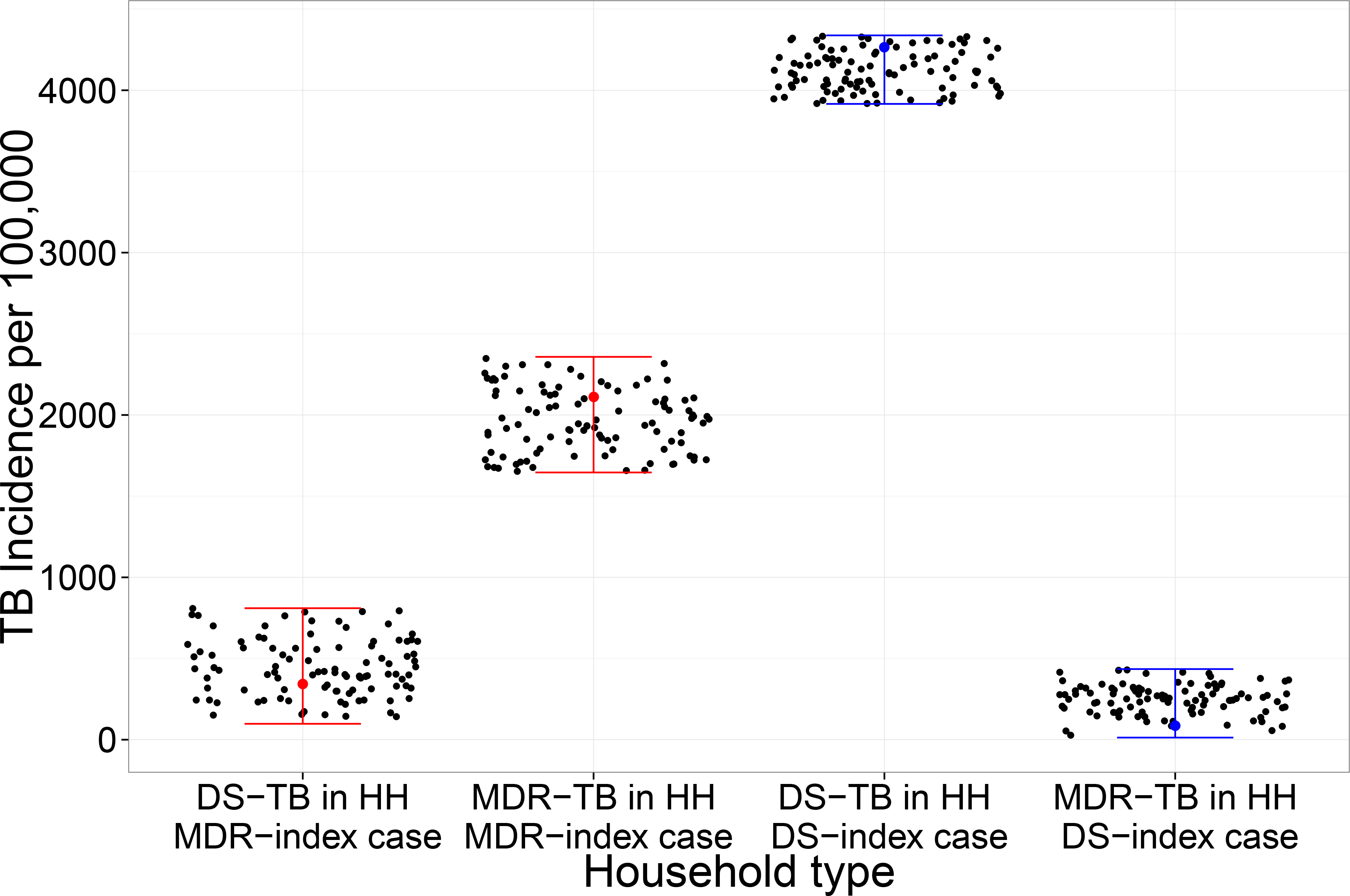
Model fits. Black dots represent Model 1 output that matches to data shown in ranges for each type of household (HH). See SI text Figures S3&S4 for equivalent plots for Model 2&3.

### Scenario analysis

A scenario analysis was used to explore the sensitivity of Model 1 results to key natural history parameters. Firstly, we changed the initial proportion of the population latently infected with MDR-*Mtb* from 2% to 10%.

A full sensitivity analysis of the parameters kept fixed in the model fits was not possible due to limitations imposed by computation time. Instead, to determine which further scenarios to explore, we determined the parameters most correlated with TB incidence in our model, and hence likely to have the biggest impact on our model fit and parameter estimates. To determine these parameters, we used LHS to choose 10,000 parameter sets from (uniform) prior distributions for all parameters (Table 2). We then ran Model 1 with these 10,000 parameter sets and determined the parameters that were statistically significantly correlated with any of the four TB incidence outputs (Kendall correlation, *p < 0.01*). These parameters were then used to design two scenarios - one with a combination of these parameters at their prior values which gave highest TB incidence and the combination which gave the lowest TB incidence.

We also increased our 10-year initial run-in period for the population to 30 years and explore the impact on the estimates. Furthermore, we explored removing the assumption of saturating household transmission (per capita transmission rate was then not dependent on household size, i.e. density-dependent transmission).

All code is available online [27].

## Results

### Fit to the data

Model structures 1-3 could all replicate the data from the household study (Figure 2). The MCMC trace and density plots of the posterior distributions are shown in S1 Text.

### Parameter estimates

The estimates of the external force of infection for DS- and MDR-TB were similar across the three models (Table 3, Figure 3). The per capita transmission rate of DS-TB within households was also similar across the three models. The relative fitness of MDR-*Mtb* was similar for Model 1 and 2, but increased in Model 3, as might be expected as in this third model the reduction in fitness is applied to two rates. For Model 1, that is assuming a resistance phenotype affects transmission, the relative fitness of MDR-*Mtb* was estimated to be 0.33 (median, 95% CI: 0.17-0.54) vs. DS-*Mtb* with a fitness of 1. In Model 2, where a resistance phenotype affected disease progression, a similar relative fitness was estimated: 0.39 (0.26-0.58). If both rates were affected, then the relative fitness of MDR-*Mtb* was estimated to be 0.57 (0.43-0.73) (Table 3, Figure 3).

**Table 3:** Parameter estimates for the median and 95% credible intervals of the four unknown parameters from at least 100 5,000 MCMC runs. The fitness cost to resistance is assumed to affect transmission in Model 1, progression to active disease in Model 2 and both transmission and progression in Model 3.

| Model | $foi_s$ | $foi_r$ | $\beta$ | $f$ |
| --- | --- | --- | --- | --- |
| 1 | 0.19 (0.04 – 0.42) | 0.08 (0.00 – 0.22) | 69.95 (53.78 – 86.86) | 0.33 (0.17 – 0.54) |
| 2 | 0.24 (0.06 – 0.46) | 0.13 (0.01 – 0.28) | 69.39 (53.46 – 86.49) | 0.39 (0.26 – 0.58) |
| 3 | 0.20 (0.05 – 0.43) | 0.11 (0.01 – 0.27) | 70.46 (54.14 – 88.19) | 0.57 (0.43 – 0.73) |

**Figure 3:**
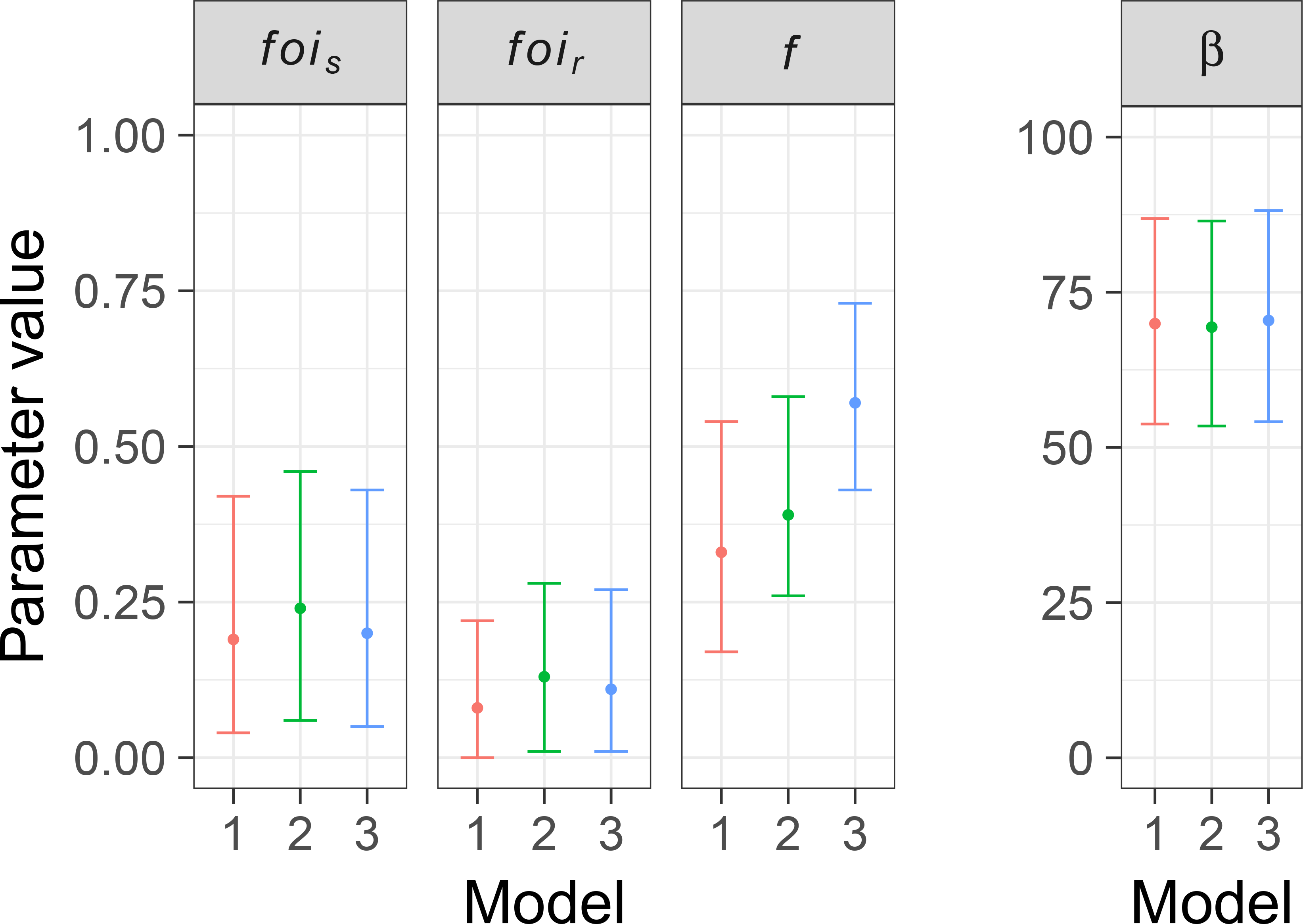
Fitted parameters from each Model. The units for the y-axis of the corresponding plots are: for the external forces of infection (‘*foi_s_*’ and ‘*foi_r_*’) proportion infected per year, for the relative fitness (*f*) there are no units and for the per capita transmission rate (‘beta’) the units are effective contact rate per year. Model 1 assumes a transmission cost to resistance, Model 2 a disease progression cost and Model 3 assumes an effect on both.

Comparing the external force of infection for DS- vs. MDR-TB we found that the ratio of the two was around 0.5 (median estimate 0.42 / 0.54 / 0.55 from the three models). This single value for the external force of infection (*foi*) represents a complex set of processes (contact patterns, length of infectiousness etc.) and so cannot be used to determine relative fitness. However, the ratio is in the range that supports our estimates of the relative fitness from the internal household model. The ratio of an approximate effective reproduction number for MDR- and DS-TB also supported our main results (see S1 Text).

### Probability of remaining free from tuberculosis

We explored the probability of remaining free from tuberculosis as was presented from the original study (Figure 2 in [17]). By comparison we had highly similar dynamics to the study (Figure 4a vs. (b-d)).

### Scenario analysis

Our five scenarios gave very similar estimates for the relative fitness of MDR-*Mtb* (a range of medians from 0.22 – 0.41, S1 text). This suggests that the estimates of relative fitness are robust to: increasing the initial proportion of households that were initially infected with latent MDR-*Mtb* from 2% to 10% (in the pre-study), setting TB incidence to high or low levels (see S1 text for parameter details), extending the initial run-in period from 10 to 30 years or removing the saturation of transmission within households.

## Discussion

Our results suggest that the average relative fitness of MDR-*Mtb* strains circulating in households in Lima, Peru in 2010-2013 was substantially lower than that of drug susceptible strains (~40-70% reduction). When a resistance phenotype was assumed to affect both transmission and progression to disease rates, then the relative fitness of MDR-TB strains was ~60%.

The strengths of this study are that we were able to fit a stochastic household-level model to detailed location-specific data, accounting for accurate distributions of both household size and study follow-up time. We were also able to differentiate between internal and external transmission, matching the resistance typing data from the household study [17]. Moreover, our transmission rate estimates account for the longer infectiousness of MDR-TB cases (due to delays in diagnosis and treatment initiation etc.). This model and its MCMC fitting algorithm can be applied to other settings and then used as the basis for predictions of future levels of DS- and MDR-TB. In particular, this novel way of estimating fitness costs, by fitting dynamic transmission models to resistance-specific incidence data could be used for other TB prevalent settings or for other bacteria. Furthermore, the estimates given can be directly translated into dynamic transmission models for prediction whilst previous estimates, for example of differences in growth rates have less clear epidemiological translations.

Our modelling analysis is limited by the assumption of homogeneity - of both hosts and strains. The characteristics of the DS- and MDR-TB contacts under consideration in the underlying household study were highly similar [17]. Thus, as our estimate is of a relative fitness we believe that including host differences in our model may have had little effect on our relative results. Strain heterogeneities however, mean that our result is (potentially) an average across many different drug resistant strains. It is known that differences in resistance and compensatory mutation combinations result in a diversity in fitness across strains [13]. This diversity is highly important for predictions of MDR-TB levels in the future [28]. Our estimate must therefore be taken as a population average in Lima, at a certain time and indicative of the mean fitness rather than an indicator of the range of potential fitness in the population. If one highly fit MDR-TB strain were to emerge (or were already present), then future prevalence predictions based on our (mean) estimate could be an underestimate. We fitted the model to data with confidence intervals that were derived without fully accounting for the dependency of infection between household members. Improving methods for robust approximation of parameters from mechanistic models that take full account of such dependencies is an important active area of research [29], and will improve future studies of this kind.

Our Model 1, where a transmission effect is assumed, is the most similar to previous models of MDR-TB transmission [6, 9, 22]. Reductions in transmission could arise from many factors including differences in location of infection (pulmonary vs. non-pulmonary), a different interaction with the basic immune system or different aerosolization levels. However, our MDR-TB fitness predictions are at the lower end of the range seen previously [10]. This may reflect the situation in Peru where there is a strong tuberculosis control infrastructure with a well-developed MDR-TB treatment program and a growing economy. These two factors may have combined to limit the spread of MDR-TB and hence prevent the adaptation of MDR-TB to a higher fitness. At the bacterial level, compensatory fitness mutations that could influence the ability of drug resistant *Mtb* strains to spread may not have emerged or not been allowed to spread. Calibrating the model to other settings would help clarify this issue. Alternatively, it may be that our estimates are providing, for the first time, a better direct translation of fitness from epidemiological data to a transmission model parameterisation.

There is a paucity of evidence for whether differences in TB disease prevalence in general are due to infection or progression to disease [30]. In particular, for resistant strains it is unclear where the effect of becoming resistant should be applied in the natural history of tuberculosis infection. Both Snider and Teixeira [31, 32] demonstrated similar levels of tuberculin skin test (TST) conversion among MDR- and DS-TB household contacts but lower levels of disease in contacts of those with MDR-TB. This was also seen in a recent study in children [33], whilst a higher prevalence of TST positivity was found in household contacts of MDR-TB patients than contacts of newly diagnosed TB patients in Viet Nam [34]. This evidence combines to suggest that the fitness cost to resistance, if any, was to be observed on the progression to disease. We make this assumption in our Model 2, where the hypothesis is that those with active TB disease, whether due to resistant or susceptible bacteria, have a similar bacterial load and hence ability to transmit successfully. However, once successfully established in a new host, resistant bacteria may be less able to combat the immune system and establish a disease state. This has been assumed in a previous model of HIV and MDR-TB interaction [35].

Previous models have assumed that resistant strains could become more fit (i.e. have a relative fitness greater than 1), whilst we capped the relative fitness of the resistant strains at 1, due to the data from previous studies and the literature [13, 36]. Our posterior parameter distributions for the estimated relative fitness parameter (reflected in the 95% CI for *f*, see S1 text) suggest that this is a valid assumption for the resistant strains circulating at this time in Lima. Importantly, all our estimates are of “relative” fitness, and therefore should be robust to changes in natural history assumptions as these would affect both drug susceptible and resistant strain transmission.

Future work will include adding in detail on host and strain heterogeneity to the model. Data collection of strain heterogeneity along with active contact tracing and an understanding of where and from whom transmission occurs would drastically improve our understanding of fitness and hence improve estimates of future MDR-TB levels. Exploring the external infection methods and potential changes in this force of infection over time (i.e. making it dynamic as in [37].) would allow for models that can predict levels of MDR-TB in Lima. Future predictive transmission modelling using our relative fitness estimates are likely to suggest that if treatment objectives are maintained and this fitness measure remains constant, that MDR-TB prevalence will remain under control in Lima in the short term.

In conclusion, we find the fitness cost of drug resistance in *Mtb* in Lima Peru to be substantial. Importantly this paper provides direct transmission model estimates, using a novel method, of the relative fitness levels of drug resistant *Mtb* strains. If these fitness levels do not change, then the existing TB control programmes are likely to keep MDR-TB prevalence at their current levels in Lima, Peru. These estimates now need to be gained for *Mtb* in other settings and the values used in models to explore future global burden.

## Author’s contributions

GMK and LG conceived of, and designed the study, with support from MZ, RHG and JF; GMK performed the mathematical modelling, with calibration support from SF. All authors contributed to analysis and interpretation. The manuscript was drafted by GMK, MZ, SF and LG, with support from JF and RHG. All authors gave final approval for publication.

## Acknowledgements

We would like to thank David Moore, Christophe Fraser and Eduardo Gushiken for their input into the initial grant. We are also grateful to anonymous reviewers from TBMAC and to the TB Modelling Group at the London School of Hygiene & Tropical Medicine for their comments. GK is affiliated with the National Institute for Health Research Health Protection Research Unit (NIHR HPRU) in Healthcare Associated Infections and Antimicrobial Resistance at Imperial College London in partnership with Public Health England (PHE). The views expressed are those of the authors and not necessarily those of the NHS, the NIHR, the Department of Health or Public Health England.

